# Dissecting the functions of NIPBL using genome editing: The importance of the N-terminus of NIPBL in transcriptional regulation

**DOI:** 10.1101/616086

**Authors:** Kosuke Izumi, Kazuhiro Akiyama, Katsunori Fujiki, Koji Masuda, Ryuichiro Nakato, Aiko Otsubo, Masashige Bando, Katsuhiko Shirahige

## Abstract

Cornelia de Lange syndrome (CdLS) is characterized by craniofacial dysmorphisms, intellectual disabilities, growth retardation, and several other systemic abnormalities. CdLS is caused by heterozygous germline mutations in structural and regulatory components of cohesin. Mutations in *NIPBL*, which encodes regulatory subunit of cohesin, are frequently found in individuals with CdLS. CdLS is associated with a currently unknown mechanism of global transcriptional dysregulation. In this study, *NIPBL* mutants were generated using the CRISPR/Cas9 system to study this mechanism. Clones with a biallelic frameshift mutation in exon 3 of *NIPBL*, resulting in a truncated N-terminus, displayed transcriptional dysregulation without sister chromatid separation defects. Detailed transcriptome analysis revealed the overexpression of genes in *NIPBL* mutants that are typically expressed at low levels in wild type and the reduced expression of genes that are typically expressed at high levels in wild type. This result suggested that NIPBL plays a role in fine-tuning gene expression levels. MAU2 protein, that closely interacts with NIPBL, was nearly absent in these clones. The reduction of MAU2 observed in *NIPBL* mutants points to the importance of the NIPBL N-terminus/MAU2 interaction in transcriptional regulatory role of NIPBL.

## INTRODUCTION

Cornelia de Lange syndrome (CdLS) is a multisystem developmental disorder characterized by facial dysmorphisms, short stature, congenital heart disease, limb defects and intellectual disability (1). CdLS is caused by genetic mutations in cohesin regulatory molecules and cohesin structural components encoded by genes including *NIPBL*, *HDAC8*, *SMC1A*, and *SMC3* (2–6). The cohesin complex functions in multiple biological processes including regulation of sister chromatid separation (SCS), DNA repair, DNA looping, and regulation of gene expression (7). The cohesin complex is made up of the structural proteins SMC1A, SMC3, RAD21 and STAG1/2 and forms a ring-like structure around DNA. NIPBL (Nibbed-B-like protein) forms a heterodimer with MAU2. The NIPBL-MAU2 heterodimer plays an important role in the loading of the cohesin complex onto DNA (8,9). Another protein, HDAC8, controls the release of the cohesin complex from DNA (6). Among the genes implicated in CdLS, mutations are most frequently identified in *NIPBL* (10).

The cohesin complex was originally identified by its ability to prevent premature SCS during mitosis, but SCS defects were not observed in cells derived from CdLS probands (11,12). Transcriptome analysis of CdLS probands revealed unique transcriptome signatures, suggesting that the molecular mechanism leading to the CdLS phenotype is caused by transcriptional dysregulation (13,14).

We used the CRSIPR/Cas9 system to introduce *NIPBL* mutations into the cell lines 293FT and HCT116 and evaluated any changes in cellular phenotypes (15). Previously, how the cohesin complex executes the multiple biological functions such as sister chromatid cohesion and transcriptional regulation, were unknown. In this study, observations of the *NIPBL* genome-edited lines showed that the N-terminus of NIPBL has a transcriptional regulatory function. Furthermore, the detailed transcriptome analysis revealed that NIPBL is involved in fine-tuning gene expression levels.

## MATERIALS AND METHODS

### Cell culture and genome editing

Cell lines 293FT and HCT116 (Courtesy of Professor Tetsu Akiyama, Laboratory of Molecular and Genetic Information, Institute of Molecular and Cellular Biosciences, The University of Tokyo, Japan) were cultured in Dulbecco’s modified Eagle medium containing 10% fetal bovine serum and 1% penicillin-streptomycin. 293FT originates from embryonic kidney cells, and HCT116 originates from colorectal carcinoma cells. The empty guide RNA (gRNA) expression vector (ID 41824) and hCas9 vector (ID 41815) were obtained from Addgene (Cambridge, MA). The gRNA target sequences GAAGAGGTGAACTGCCTTT (*NIPBL* exon 3) and CTCGTTCTGATTTTAACCG (*NIPBL* exon 10) were cloned into the gRNA empty vector using Phusion polymerase (M0530S: New England Biolabs, Ipswich, MA) and the Gibson assembly system (E5510S: New England Biolabs). Cas9 and gRNA vectors were transfected into the target cell lines using Lipofectamine 2000 transfection reagent (Thermo Fisher Scientific, Waltham, MA) and the Neon electroporation transfection system (Thermo Fisher Scientific). Genomic DNA was extracted using the NucleoSpin tissue kit (Macherey-Nagel, Duren, Germany) and sequenced by Sanger sequencing. Primers for *NIPBL* exons 3 and 10 are available upon request. PCR products, pMD20 vector, and NEB10 beta competent cells were used to characterize the 293FT ex10mut mutant cell line by TA cloning.

### Western blotting

SDS sample buffer was added to transfected cells to obtain total cellular lysates, which were purified by chromatin fractionation (14). Obtained lysates were loaded onto Mini-PROTEAN TGX Precast gels (product #456-1086, Bio-Rad, Hercules, CA) and transferred onto a nitrocellulose membrane for western blotting. Antibodies were directed against the following: NIPBL N-terminus (sc-374625, Santa Cruz Biotechnology, Dallas, TX); NIPBL exon 10 (A301-779A, Bethyl laboratories Montgomery, TX); NIPBL C-terminus (AM20797 KT55, Acris Antibodies, Rockville, MD); SMC1A (ab21583, Abcam, Cambridge, MA); MAU2 (SCC4; ab469069, Abcam, Cambridge, MA), and histone H3 (ab1791; Abcam, Cambridge, MA). Antibody to α-tubulin were (T6074) from Sigma-Aldrich. Two biologically independent replicates were performed.

### Giemsa staining and nuclear area measurements

Nocodazole (100 ng/ml; Calbiochem, Merck Millipore, Darmstadt, Germany) was used to treat 293FT and HCT116 cells for 3 h. Treated cells were allowed to swell in a hypotonic solution of dilute phosphate-buffered saline for 5 min at room temperature and then fixed with Carnoy’s solution prior to being dropped onto glass slides. The slides were stained with Giemsa solution (Merck, Billerica, MA), washed with water, and then mounted with Entellan water-free mounting medium (Merck). The presence or absence of SCS was evaluated by manual inspection of 200 293FT cells and 150 HCT116 cells. Nuclear area was measured using fluorescence *in situ* hybridization (16). Briefly, cells were fixed with a hypotonic solution comprising 0.075 M KCl, EGTA, Hepes, and a fixative solution (methanol:glacial acetic acid, 3:1 v/v). DAPI stain was used to stain nuclei for size measurement using an Olympus BX51 microscope and MetaMorph software (Molecular Devices, Sunnyvale, CA). The nuclear area of ~150 cells was measured for each sample (159 WT cells, 146 ex3mut cells and 173 ex10mut cells).

### Fluorescence-activated cell sorting

Ethanol, followed by a phosphate-buffered saline wash, was used to fix cells. DNA was stained with a propidium iodine solution, and DNA profiles were evaluated using fluorescence-activated cell sorting with a FACSCalibur flow cytometer and Cell Quest software (BD Biosciences, San Jose, CA).

### RNA sequencing

Total RNA was extracted by TRIzol (Life Technologies, Carlsbad, CA) and NucleoSpin RNA (Macherey-Nagel, Duren, Germany). All samples were sequenced on the Illumina HiSeq 2500 platform as single-end 50-bp reads and mapped to the human genome (UCSC build hg19). Sequencing and mapping statistics are summarized in Supplementary Table S1. The RNA sequencing analysis (RNA-seq) was performed with TopHat and Cufflinks (17,18). Only the protein-coding genes were considered for the transcriptome analysis, and gene-level expression values were estimated by reads per kilobase of exon per million fragments mapped (RPKM) score. Translational start sites prediction was performed with NetStart 1.0 translation initiation site prediction program (http://www.cbs.dtu.dk/services/NetStart/)(19).

### DRB-Nascent RNA sequencing

Before capturing nascent RNA from 293FT cells, 50 µM 5,6-dichloro-1-β-D-ribofuranosyl benzimidazole (DRB) was added to the culture medium to inhibit transcription for 30 min and align the transcription progress. Afterwards, the medium was replaced with DRB-free medium, and 0.5 mM ethylene uridine was added to the medium with subsequent incubation for 60 min. Total RNA was prepared with TRIzol reagent (Invitrogen). A total of 1 µg of ethylene uridine–labeled RNA was biotinylated with 0.5 mM biotin azide using the Click-iT Nascent RNA Capture kit, in accordance with the manufacturer’s instructions (Thermo Fisher Scientific, Waltham, MA).

Biotinylated RNA was precipitated with chilled ethanol and resuspended in Ultra-Pure™ DNase/RNase-free distilled water. The purified biotinylated RNA was mixed with Dynabeads MyOne Streptavidin T1 magnetic beads in Click-iT RNA binding buffer and heated at 70°C for 5 min, followed by incubation at room temperature for 30 min with gentle agitation. Following washing with Click-iT wash buffer 1 and wash buffer 2, the beads were resuspended in wash buffer 2 and used for cDNA synthesis. cDNA was synthesized using the SuperScript® VILO™ cDNA synthesis kit (Thermo Fisher Scientific) in accordance with the manufacturer’s instructions. cDNA was amplified using a slightly modified in vitro transcription amplification method (20). Second-strand cDNA synthesis was used to assemble a library and sequenced following the manufacturer’s instructions (NEB Next DNA Library Prep Master Mix Set for Illumina; NEB E6040L). Sequenced reads were aligned to the human reference genome hg19 using TopHat v2.1.0 with the ‘--no-coverage-search --read-realign-edit-dist 0’ option selected (17). Averaged nascent DNA strand profiles were calculated and visualized using custom scripts written in Python and R. Information and statistics regarding sequenced reads and mapping rates are summarized in the Supplementary Table S1.

### ChIP-seq

Both 293FT wild-type (WT) and *NIPBL* genome-edited cell line clones were used for ChIP-seq (chromatin immunoprecipitation followed by massively parallel sequencing) (6,14) with a custom-made antibody against RAD21(6,14). Harvested input and ChIP DNA samples were sequenced by the Illumina HiSeq 2500 Sequencing System (San Diego, CA). Sequenced reads were aligned to the human reference genome hg19 using Bowtie software set to allow two mismatches in the first 28 bases per read and output only uniquely mapped reads (-n2 –m1 option). Mapping statistics are summarized in Supplementary Table S3. DROMPA v2.5.3 software was used for ChIP-seq and visualization (21). To facilitate comparison of detected peaks between ChIP experiments, the number of mapped reads per cycle was normalized to the total number of mapped reads. The average reads plot around a transcription start site (TSS) was obtained using the PROFILE command, and RefSeq gene annotation was obtained from the UCSC genome browser (http://genome.ucsc.edu/).

## RESULTS

### CRISPR genome editing of the *NIPBL* locus

The presence of a haploinsufficient *NIPBL* with frameshift mutations in exons 3 and 10 has been observed in patients with CdLS (21). gRNAs targeting exons 3 and 10 were designed to model genetic alterations seen in patients with CdLS (Figure 1A). Exon 10, encoding a coiled-coil region and an undecapeptide repeat, is a CdLS mutation hotspot (10). We introduced *NIPBL* mutations into two commonly used human cell lines, 293FT and HCT116. Mutations were introduced via transfection of Cas9 and gRNA plasmids into cells, and single-cell cloning was performed using the limiting dilution technique. The presence of *NIPBL* mutations was confirmed by Sanger sequencing.

**Figure 1.**
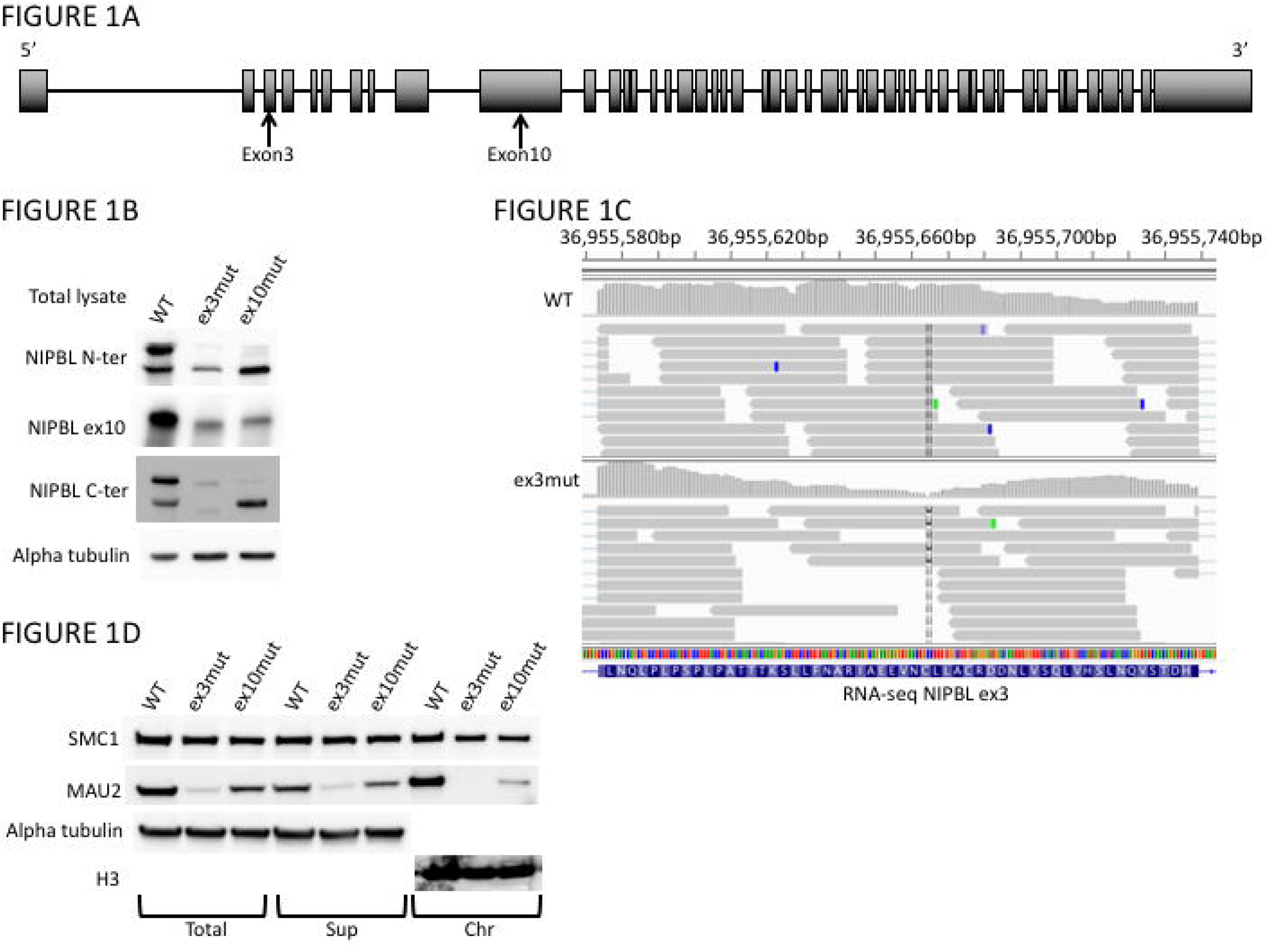
293FT *NIPBL* mutant cell lines. (**A**) *NIPBL* gene structure. Exon and intron lengths are not proportional. gRNA target sites within exons 3 and 10 are indicated by arrows. (**B**) Western blotting results for NIPBL. Use of an antibody against the N-terminus (N-ter) or against the C-terminus (C-ter) yielded two bands. The upper band represents full-length NIPBL, and the lower band represents the shorter isoform. Western blotting with clone ex3mut total cellular lysate and the N-ter antibody demonstrated the absence of full-length NIPBL. Western blotting with the C-ter antibody demonstrated the presence of truncated NIPBL. Western blotting of clone ex10mut lysates demonstrated a decreased amount of full-length NIPBL and an increased amount of the truncated NIPBL isoform. (**C**) RNA-seq results for the *NIPBL* locus. The IGV browser view shows the presence of the *NIPBL* transcript in clone ex3mut. Transcripts lacked only a few base pairs targeted by *NIPBL* gRNA. (**D**) Western blotting for MAU2 and SMC1A in the total cellular lysate, soluble fraction (Sup), and chromatin fraction (Chr) in 293FT cell line clones. MAU2 was almost absent in the chromatin fraction of clone ex3mut, and its level was reduced in clone ex10mut.

A total of 20 clones with *NIPBL* exon 3 mutations in 293FT cells were screened. One clone, designated ex3mut, was identified to have 4-bp and 1-bp deletions (Supplementary Figure S1A). A total of 18 clones were screened with *NIPBL* exon 10 mutations in 293FT cells. One clone, designated ex10mut, was identified to have two frameshift mutations in exon 10 as well as an intact NIPBL allele (Supplementary Figure 1B). Sanger sequencing of clone ex10mut revealed the presence of more than three copies of *NIPBL*. Although the 293FT cell line did not originate from cancerous tissue, it is well known that 293FT-derived cell lines exhibit aneuploidy, suggesting the presence of more than two alleles of *NIPBL* in 293FT cells (22).

Karyotyping has revealed HCT116 cell lines to be generally diploid, with polyploids reported to occur at a rate of only 6.8% (http://www.atcc.org/products/all/CCL-247.aspx#characteristics). A total of 43 clones were screened with *NIPBL* exon 3 mutations in HCT116 cells. One clone, designated ex3mut1, was identified to have biallelic frameshift mutations (Supplementary Figure S1C). Another clone, designated ex3mut2, was identified to have frameshift mutations and a single intact *NIPBL* allele (Supplementary Figure S1D). A total of 21 clones were screened with *NIPBL* exon 10 mutations in HCT116 cells, and none had biallelic loss-of-function mutations in *NIPBL*. Two clones, designated ex10mut1 and ex10mut2, were obtained with only in-frame deletion/frameshift mutations (Supplementary Figure S1E and S1F). The clones described above were isolated and used for all subsequent experiments (Table 1).

**Table 1.**
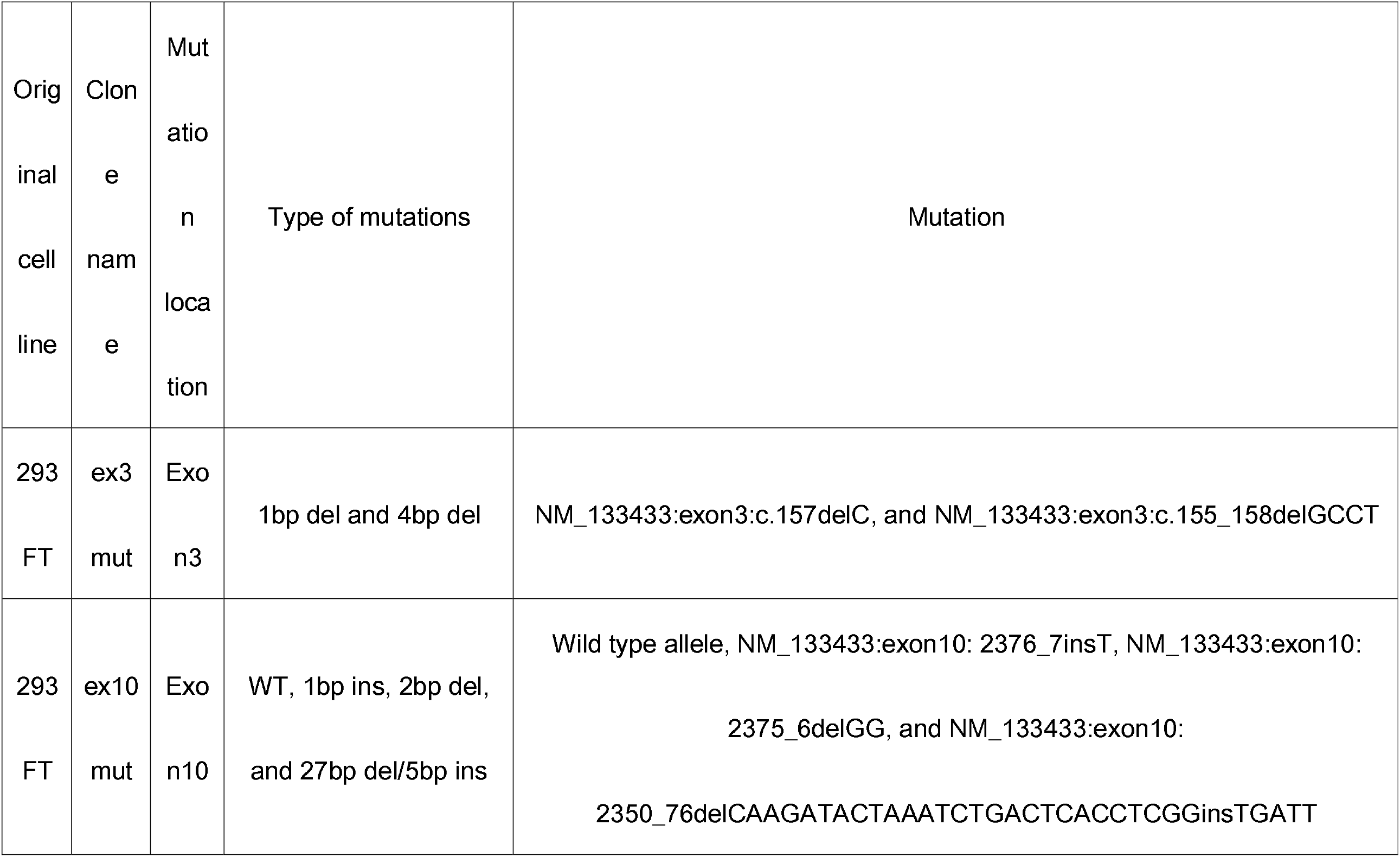
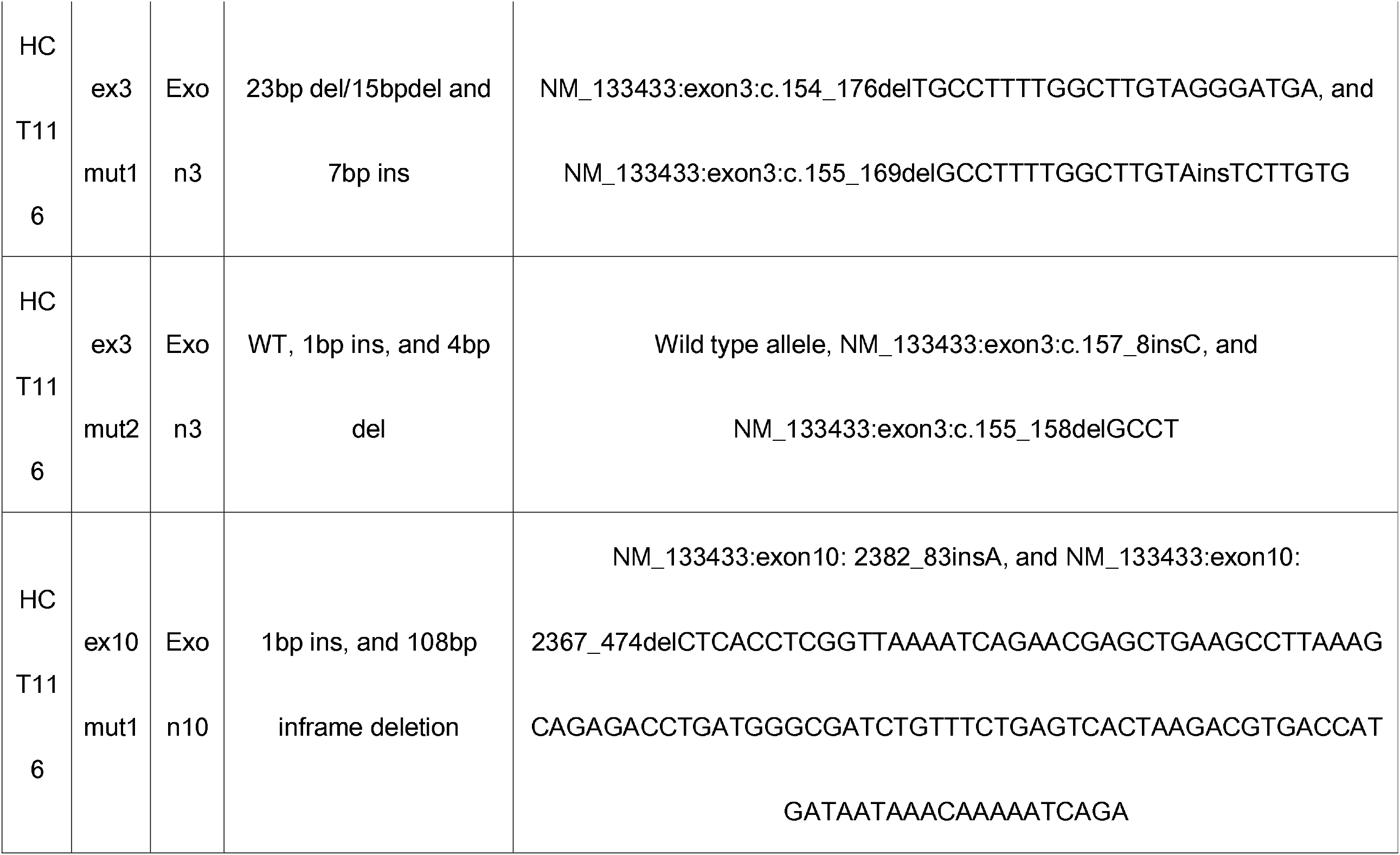
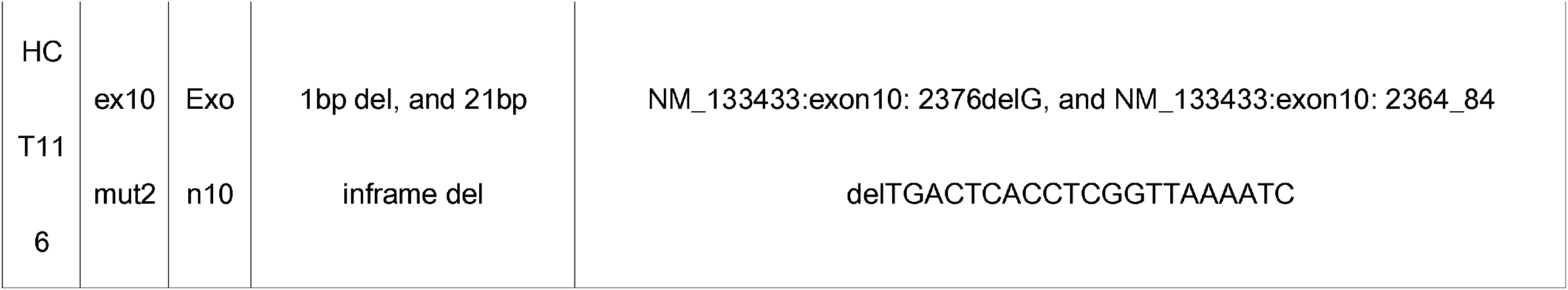
Characteristics of *NIPBL* mutant clones in cell lines 293FT and HCT116.

### Consequences of *NIPBL* mutations

To evaluate the effect of frameshift mutations in *NIPBL*, we performed western blotting using NIPBL antibodies in CRISPR genome-edited cell lines. Three different NIPBL antibodies were used, including one that recognizes the NIPBL N-terminus (amino acid residues 1–300; sc-374625), one that recognizes the protein-coding region of exon 10 (A301-779A), and one that recognizes the C-terminus of NIPBL short form (isoform B; AM20797 KT55). Western blotting of WT 293FT cells yielded two main bands with both NIPBL N-terminus and C-terminus antibodies and a single band with the exon 10 antibody, which corresponded to the molecular size of the upper band of the former set of two bands (Figure 1B). The two bands resulting from the N-terminus and C-terminus antibodies represent the full-length NIPBL and an isoform of NIPBL lacking exon 10, respectively. A similar NIPBL isoform was recently reported (23). Results from reverse transcription-PCR analysis of WT, clone ex3mut, and clone ex10mut, revealed the presence of a NIPBL isoform lacking exon 10 (Supplemental Figure S3).

Western blotting of 293FT cell–derived clone ex3mut with the N-terminus antibody revealed the absence of the top band and a decreased intensity of the lower band (Figure 1B). Additionally, use of the C-terminus antibody revealed the presence of two bands with slightly smaller molecular sizes, indicating the presence of a N-terminally truncated NIPBL (Figure 1B). Western blotting of clone ex10mut, produced from 293FT cells, showed an overall reduction in the amount of full-length NIPBL and a slight increase in the amount of the NIPBL isoform lacking exon 10, as demonstrated by the stronger intensity of the lower band (Figure 1B). Western blotting of HCT116 cell lines encoding *NIPBL* mutants produced similar results to those observed for the *NIPBL* mutant 293FT lines (Supplementary Figure S2A).

The introduction of a frameshift mutation in *NIPBL* exon 3 may activate the alternative translation initiation site located downstream of exon 3 in the gRNA target region. RNA-seq and quantitative reverse transcription-PCR revealed the presence of a *NIPBL* exon 3 transcript, supporting the potential for an alternative initiation site (Figure 1C). In exon 5, several ATG codons exist in frame. The distance between the ATG initiation codon at exon 2 and these ATG sequences in exon 5 is ~363-bp and encodes ~121 amino acid residues (~13.5 kDa). The sequence upstream of this ATG rich region in exon 5 was similarly predicted to be a translation initiation codon (NetStart initiation site prediction score = 0.454) to the initiation codon for full-length NIPBL located in exon 2 (NetStart initiation site prediction score = 0.508) (Supplementary Figure S4) (24).

### Effects of *NIPBL* deletions on cohesin and loader complexes

To evaluate the effect of the absence of full-length NIPBL, we performed western blotting with MAU2 and SMC1A antibodies following chromatin fraction separation of cell lines 293FT and HCT116. Surprisingly, the quantity of MAU2 in 293FT cell clone ex3mut and HCT116 cell clone ex3mut1 was markedly diminished in both the total cellular fraction and the chromatin fraction (Figure 1D and Supplementary Figure 1D). However, the amount of SMC1A was only slightly reduced in these cell-line clones. In 293FT clone ex10mut and HCT116 clones ex3mut2 and ex10mut1, the amount of MAU2 was slightly decreased in the total cellular lysate and decreased in the chromatin fraction (Figure 1D and Supplementary Figure S1D). RNA-seq revealed that *MAU2* expression was comparable between the control (10.9 RPKM) and 293FT clone ex3mut (12.9 RPKM); therefore, the reduction of MAU2 was likely not due to reduced gene expression.

### Effect on SCS

Because NIPBL is a cohesin loading factor and the main function of the cohesin complex is to maintain sister chromatid cohesion, the effect of *NIPBL* mutations on SCS was evaluated. The observed frequency of the SCS defects was very low in both 293FT cell clone ex3mut and HCT116 clone ex3mut1. The frequency of SCS defects in these clones was comparable to that of WT 293FT and HCT116 cells (Table 2 and Figure 2A). Normal SCS defect frequency was observed in the remaining HCT116 clones, including ex3mut2 and ex10mut1. However, the frequency of SCS defects was higher in 293FT clone ex10mut, which had a reduced amount of full-length NIPBL, including an increased amount of the NIPBL isoform lacking exon 10. This result suggested that reducing the amount of NIPBL (inclusive of residues encoded by exon 10) may have a major effect on the frequency of SCS defects.

**Table 2.** Frequency of SCS defects.

|  |  |
| --- | --- |
| 293FT | % |
| WT | 5.0 |
| ex3mut | 7.5 |
| ex10mut | 38.0 |
| HCT116 | % |
| WT | 9.3 |
| ex3mut1 | 6.0 |
| ex3mut2 | 7.3 |
| ex10mut1 | 8.7 |

**Figure 2.**
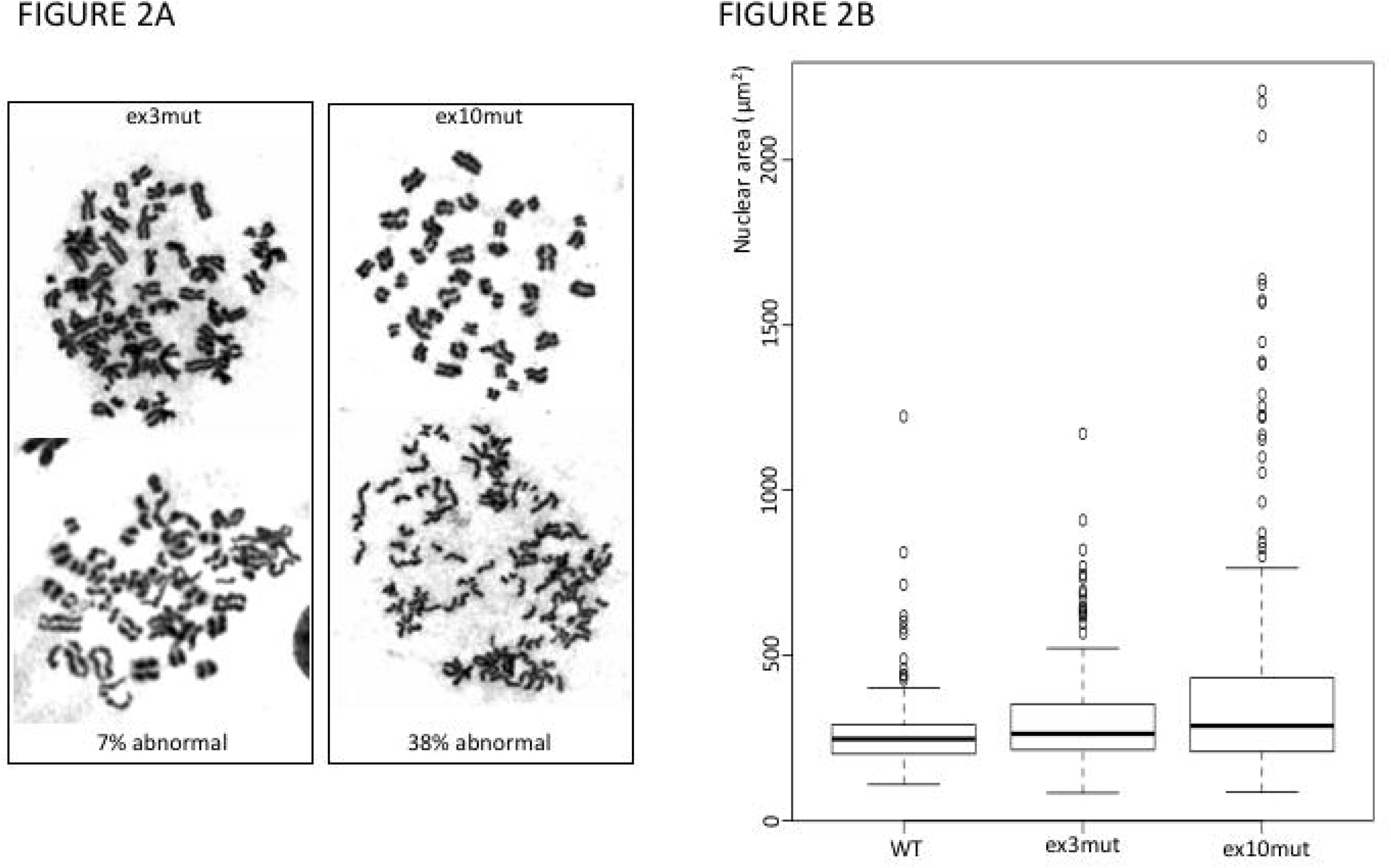
Consequences of *NIPBL* mutations. (**A**) SCS defects in targeted exon 10 mutant clones. The frequency of SCS defects was not increased in clone ex3mut, but it was increased in clone ex10mut. (**B**) Box plot showing nuclear size. Nuclear size was greater in clone ex10mut compared with the control (t-test: p-value = 1.744e-06). The y axis represents the area of the nucleus (µm^2^).

### Effect on nuclear area

Cohesin plays an important role in organizing nucleosome architecture. In cohesin component mutants of CdLS cell lines, chromatin decompaction has been observed (24). Evaluation of nuclear area in *NIPBL* CRISPR genome-edited cell lines revealed no major difference from WT in the nuclear area of 293FT clone ex3mut, but the nucleus was slightly larger in 293FT clone ex10mut (Figure 2B). Analysis of cell cycle distribution by fluorescence-activated cell sorting showed no major difference among WT 293FT cells and clones ex3mut and ex10mut (Supplementary Figure S5).

### Transcriptome analysis

We next evaluated the transcriptomic effects of *NIPBL* frameshift mutations. RNA-seq of WT 293FT cells and clones ex3mut and ex10mut revealed that both clones had similar transcriptome alteration compared to WT (Supplementary Table S1 and Figure 3[A]). The top 250 upregulated genes and 250 downregulated genes were similarly misexpressed in both clones ex3mut and ex10mut (Figure 3B, C). These observations suggested that the transcriptome effects of exon 3 and exon 10 deletions are similar. RNA-seq analysis was performed, in duplicate, for both WT 293FT cells and clone ex3mut to further define differentially expressed genes. The results showed that 485 genes were upregulated and 190 genes were downregulated, at statistically significant levels, in clone ex3mut replicates compared with WT (Supplementary Table S2).

**Figure 3.**
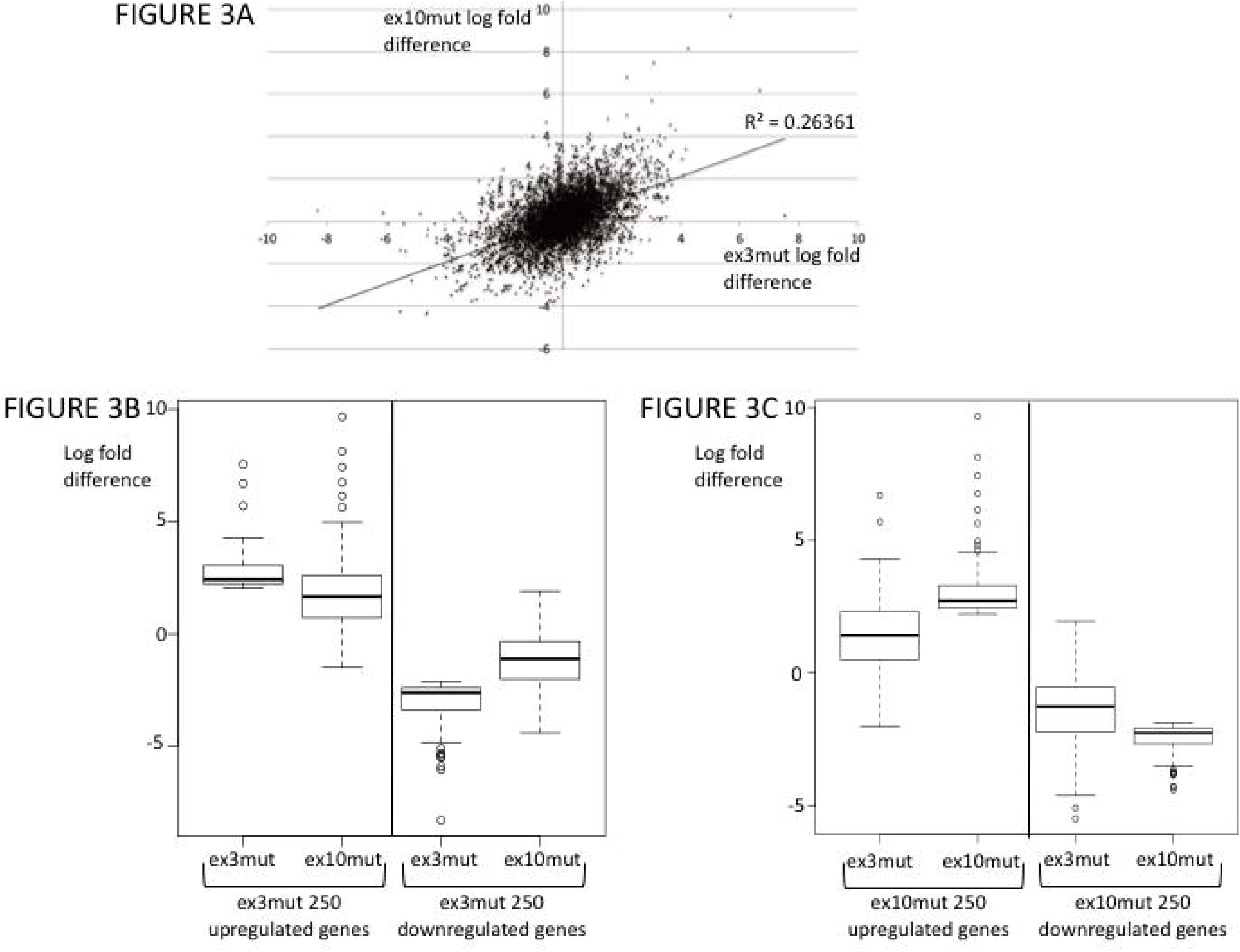
Transcriptome analysis of *NIPBL* genome-edited clones. (**A**) Scatter plot demonstrating the general positive correlation between gene expression in clones ex3mut and ex10mut. The *x* axis represents the log fold expression difference of clone ex3mut, and the *y* axis represents the log fold expression difference of clone ex10mut. *y* = 0.5062*x* + 0.0676. R² = 0.26361. (**B**) Box plot demonstrating that the differentially expressed genes in clone ex3mut were similarly misexpressed in clone ex10mut. The *y* axis represents the log fold gene expression difference observed for clone ex3mut or clone ex10mut compared with WT 293FT cells. (**C**) Box plot demonstrating that the differentially expressed genes in clone ex10mut were similarly misexpressed in clone ex3mut. The *y* axis represents the log fold gene expression difference observed for clone ex3mut or clone ex10mut compared with WT 293FT cells.

To further characterize transcriptome alterations resulting from mutations in *NIPBL*, DRB-nascent RNA-seq was performed, in duplicate, on WT 293FT cells and clone ex3mut. DRB-nascent RNA-seq is a method to quantify the amount of newly transcribed mRNA molecules within a certain period (25). The results showed a correlation of global gene transcription patterns and gene expression levels between total RNA-seq and DRB-nascent RNA-seq (R-value: 0.30; Supplementary Figure S6). Global DRB-nascent RNA-seq profile patterns were similar between WT and ex3mut clone replicates (Figure 4A). DRB-nascent RNA-seq profiles of differentially expressed genes in ex3mut showed no major differences in downregulated genes. Only the reduction of nascent RNA from the latter half of the gene coding region was observed (Figure 4C). Differences between WT and ex3mut were more apparent in ex3mut upregulated genes; increased amounts of nascent RNA transcript were seen for the entire coding region (Figure 4B).

**Figure 4.**
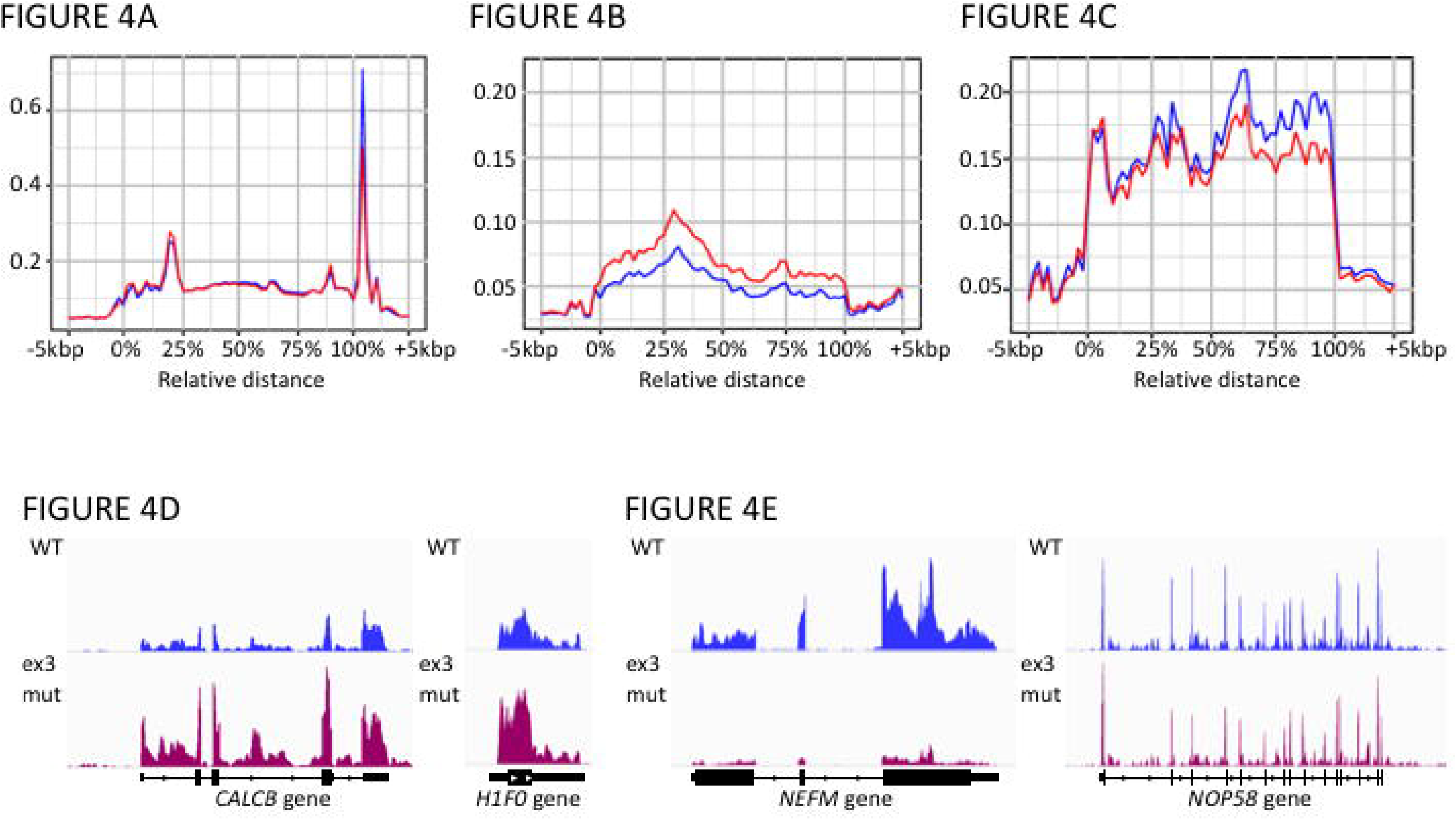
DRB-nascent RNA-seq. The read profile around the gene coding region is depicted in which the *x* axis represents the relative position of the gene. The TSS is indicated by 0%, and 100% indicates the transcription termination site. The *y* axis represents the normalized nascent strand reads. The control profile is drawn in blue, and the profile for clone ex3mut is drawn in red. (**A**) No major differences in transcription pattern were detected between WT and clone ex3mut replicates. (**B**) The transcript pattern for clone ex3mut upregulated genes (485 genes total) shows an increase in the number of transcripts throughout the gene coding region. (**C**) The transcript pattern for clone ex3mut downregulated genes (190 genes total) indicates no major differences from WT until the mid-gene coding region. (**D**) Representative loci for the ex3mut upregulated genes *CALCB* and *H1F0*. The top lane shows the read counts for the control sample, and the middle lane shows clone ex3mut sample read counts. (**E**) Representative loci for the ex3mut downregulated genes *NEFM* and *NOP58*. The top lane shows the read counts of the control sample, and the middle lane shows clone ex3mut sample read counts.

The DRB-nascent RNA-seq profile was distinctively different between ex3mut upregulated and downregulated genes. In ex3mut upregulated genes, there were no peaks around the TSS; for ex3mut downregulated genes, however, there were peaks around the TSS, which may represent nascent RNA produced from paused RNA polymerase II (Figure 4). Furthermore, comparison of nascent RNA-seq profiles between upregulated and downregulated genes indicated that expression of the downregulated genes is higher than that of the upregulated genes, suggesting that the effect of *NIPBL* deficiency is distinctively different for genes that are normally expressed at high levels compared with those expressed at low levels (Figure 5A). A similar trend was seen in HCT116 cell mutant ex3mut1 (Figure 5B). These observations prompted us to re-evaluate the RNA-seq results for a CdLS skin fibroblast sample, and indeed the gene expression pattern was similar to that observed in ex3mut mutants (14) (Figure 5C) These findings indicated that gene expression patterns are less fine-tuned in cells with *NIPBL* mutations.

**Figure 5.**
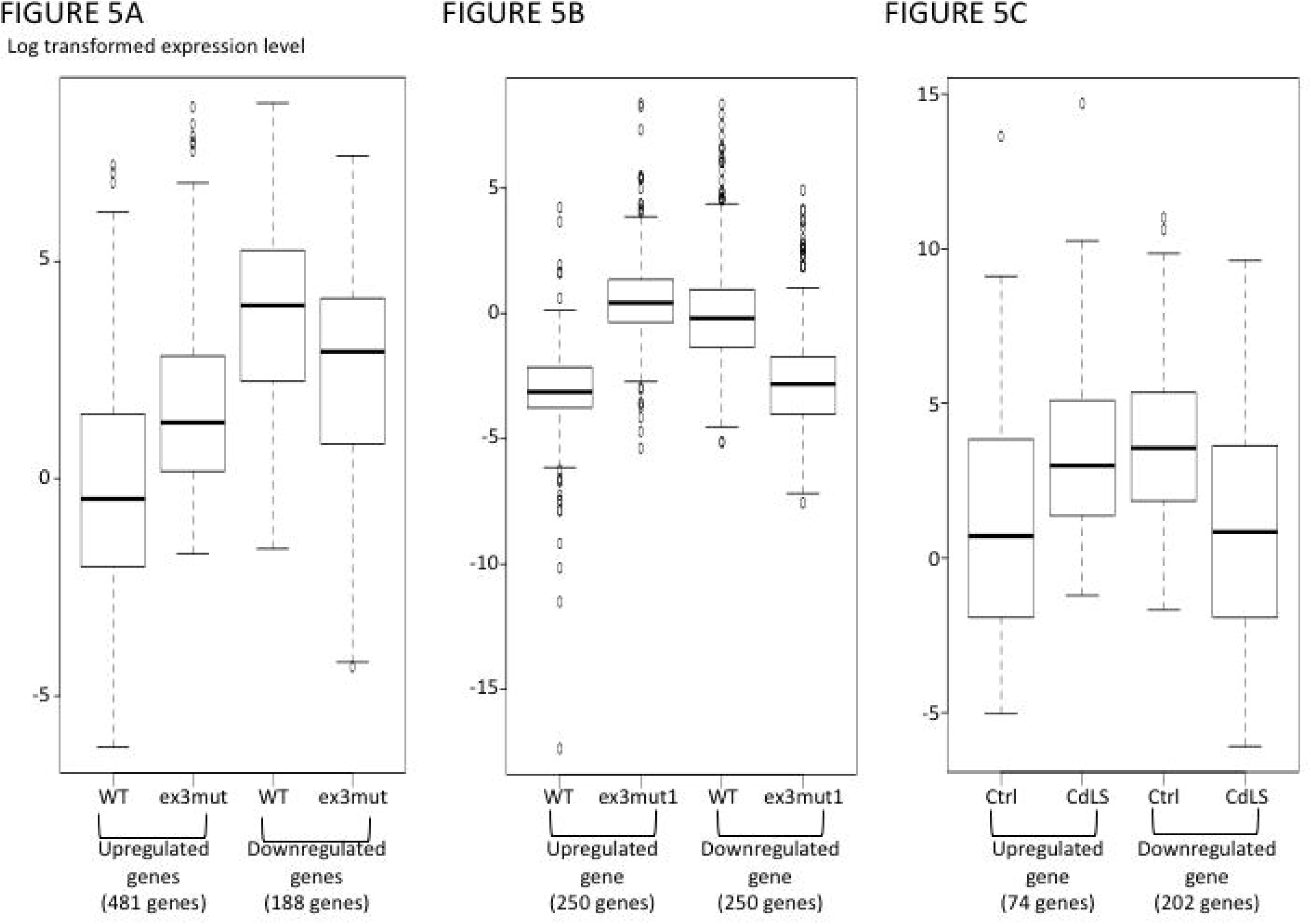
Characteristics of upregulated genes and downregulated genes in *NIPBL* mutant cell lines. The *y* axis represents the log transformed RPKM. Genes expressed below detection level in WT or mutant clones were not included in this analysis. (**A**) 293FT cell line gene expression levels for clone ex3mut upregulated genes (481 genes total) were lower than those of downregulated genes (188 genes total) in WT 293FT cells. (**B**) HCT116 cell line gene expression levels for clone ex3mut1 upregulated genes (250 genes total) were lower than those of downregulated genes (250 genes total) in WT HCT116 cells. (**C**) Gene expression analysis of a cell line from a CdLS patient revealed 74 upregulated genes and 202 downregulated genes compared with control skin fibroblast cell lines.

### RAD21 ChIP-seq

ChIP-seq with RAD21 antibody was carried out on WT 293FT cells and clones ex3mut and ex10mut to evaluate the genome-wide distribution of cohesin (Supplementary Table S3). RAD21 is a component of the cohesin complex. The number of identified RAD21 binding sites was comparable among the three 293FT lines (Table 3). The most notable difference was in peak number located upstream of genes; a ~10% reduction in peak number was observed in both clones ex3mut and ex10mut compared with WT.

**Table 3.**
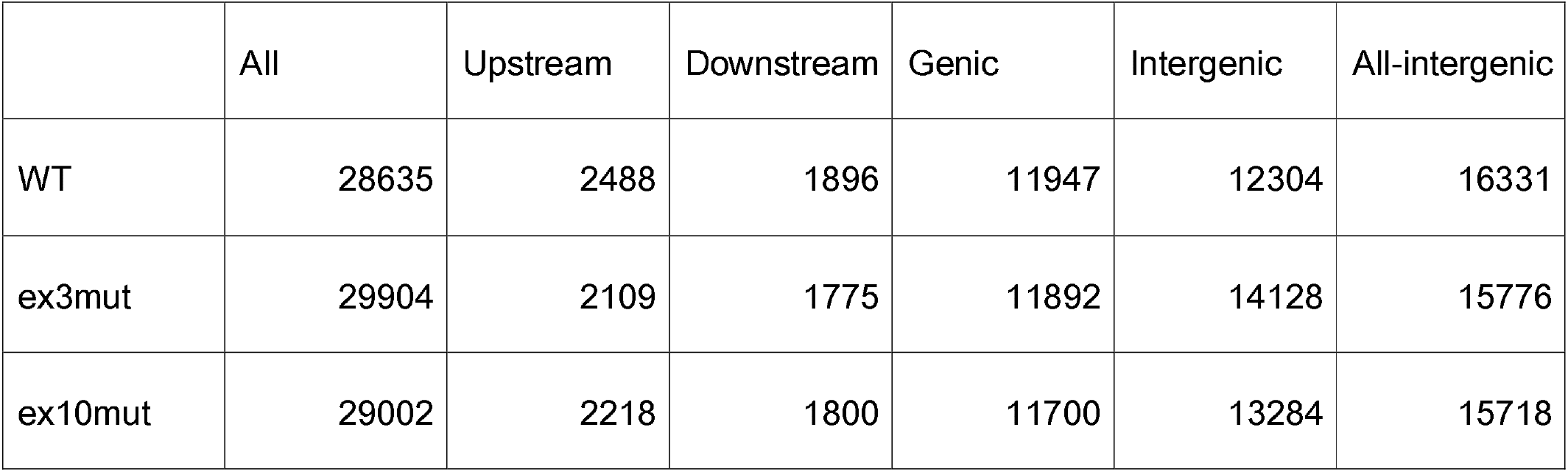
Number of identified RAD21 binding sites.

|  | All | Upstream | Downstream | Genic | Intergenic | All-intergenic |
| --- | --- | --- | --- | --- | --- | --- |
| WT | 28635 | 2488 | 1896 | 11947 | 12304 | 16331 |
| ex3mut | 29904 | 2109 | 1775 | 11892 | 14128 | 15776 |
| ex10mut | 29002 | 2218 | 1800 | 11700 | 13284 | 15718 |

TSS profile analysis was performed to further evaluate genome-wide alterations in the distribution of RAD21. The results revealed a reduction in TSS peaks in both clones ex3mut and ex10mut. These data were consistent with the observed reduction in gene upstream peak number (Figure 6A). TSS profile analysis of differentially expressed genes showed a similar reduction in RAD21 binding in both upregulated and downregulated genes (Figure 6B and 6C).

**Figure 6.**
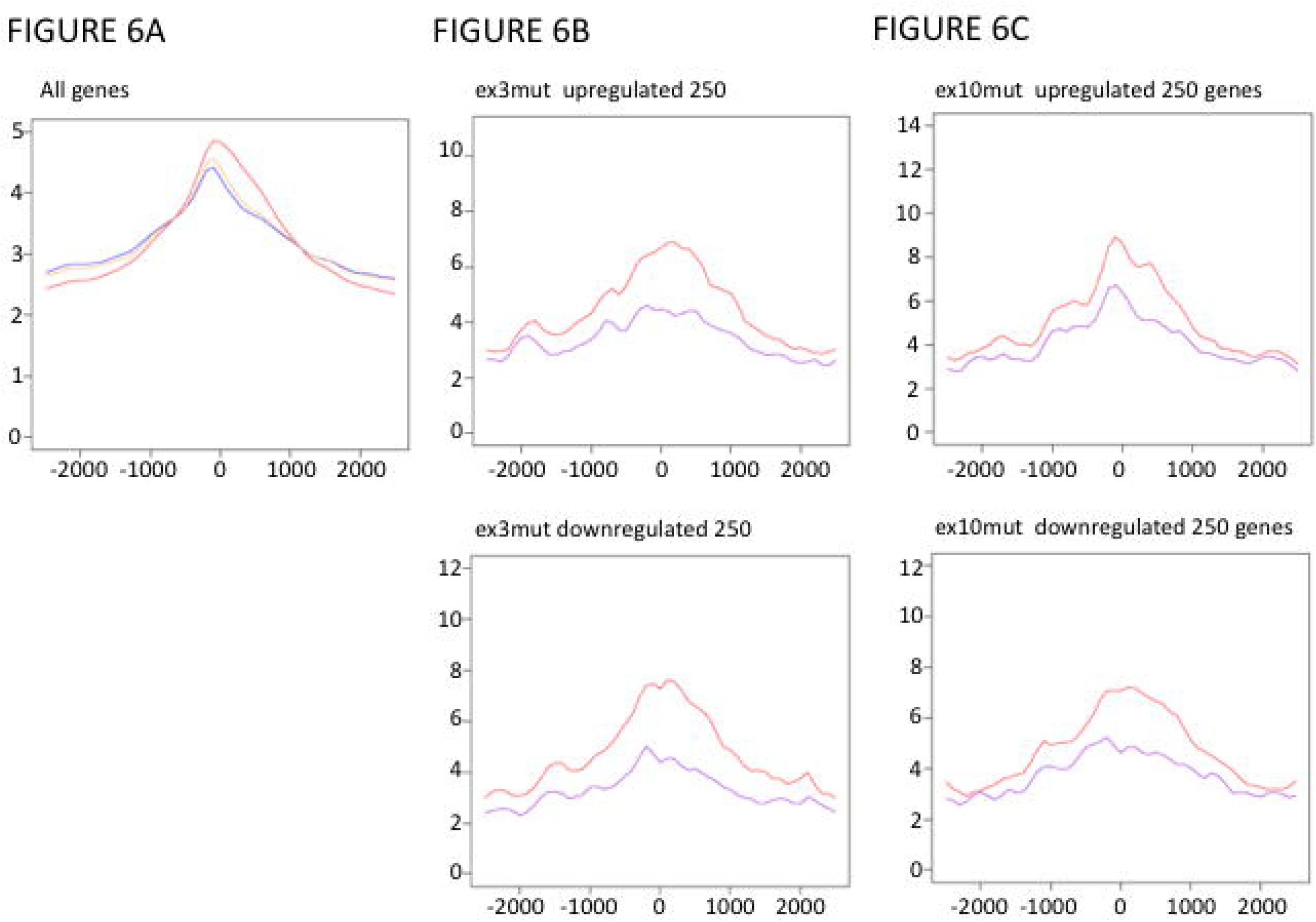
RAD21 ChIP-seq results. TSS profiles are shown in which the red line indicates the WT profile, the blue line indicates the profile for clone ex3mut, and the yellow line indicates the profile for clone ex10mut. The *x* axis represents the distance from TSS (bp), and the *y* axis represents the read density. (**A**) TSS profiles of all genes revealed the global reduction of RAD21 binding around TSS. (**B**) TSS profiles of differentially expressed genes in clone ex3mut compared with WT, in which the red line indicates the WT profile, and the purple line indicates the clone ex3mut profile. Upregulated and downregulated genes detected in clone ex3mut (top 250 genes each) indicated a similar reduction in RAD21 binding around the TSS. (**C**) TSS profiles of differentially expressed genes for clone ex10mut, in which the red line indicates the WT profile, and the purple line indicates the clone ex10mut profile. Upregulated and downregulated genes detected in clone ex10mut (top 250 genes each) indicated a similar reduction in RAD21 binding around the TSS.

## DISCUSSION

Cohesin and its loader complex have various roles in multiple biological processes, but how they carry out these roles remains unknown. Previous studies have revealed that cohesin loaders are essential for embryogenesis because *NIPBL* or *MAU2* null mice could not be obtained (26). *NIPBL* genome-edited cell lines provided a unique opportunity to evaluate the effect of the complete elimination of full-length NIPBL. Our observations indicate that full-length NIPBL is not essential for cell viability and that an N-terminally truncated NIPBL can compensate for the lack of full-length NIPBL. Based on observations of shared transcriptome alterations between *NIPBL* exon 3–targeted and exon 10–targeted clones and the absence of a SCS defect phenotype in the exon 3–targeted clone, we concluded that the N-terminus of NIPBL is mainly required for transcriptional regulation and is not necessary for the control of sister chromatid cohesion or nuclear compaction. To our knowledge, this is the first report of the functional dissection of NIPBL.

In CdLS patients, frameshift mutations in *NIPBL* exon 3 have been reported (27) that are similar to those we detected in 293FT clone ex3mut. We demonstrated that this mutation does not affect SCS. Therefore, abnormal SCS is unlikely to be the underlying disease mechanism of CdLS; rather, our data suggest that CdLS is a disorder of transcriptional regulation (28). Clone ex3mut lacks the N-terminal region of NIPBL, and the amount of truncated NIPBL was reduced; Hence, the ex3mut cells were able to tolerate a moderate reduction of NIPBL and retain normal SCS. This observation is consistent with the absence of SCS defects detected in CdLS (11,12). Furthermore, it is unlikely that MAU2 plays a role in SCS, i.e., given the absence of MAU2 in clone ex3mut and HCT116 clone ex3mut1.

Because CdLS is caused by various mutations in *NIPBL*, including whole-gene deletion, haploinsufficiency of *NIPBL* represents the underlying genetic mechanism of CdLS. Although a reduction of truncated NIPBL was observed in 293FT clone ex3mut compared with WT, the amount of truncated NIPBL in HCT116 clone ex3mut1 was comparable to WT. However, clone ex3mut1 had transcriptome abnormalities characterized by less fine-tuning of gene expression patterns. This observation indicated the importance of the NIPBL N-terminus in transcriptional regulation. The importance of the NIPBL N-terminal region, which interacts with MAU2, is further supported by the presence of CdLS probands harboring missense mutations in this region. *NIPBL* missense mutations (G15R and P29Q) that alter MAU2 binding to NIPBL were identified in individuals with CdLS (29). The same study demonstrated that the 38-residue amino acid sequence of the N-terminus of NIPBL is important for MAU2 binding (29). This sequence is likely missing in 293FT clone ex3mut and HCT116 clone ex3mut1. *NIPBL* exon 3– and exon 10–targeted clones both had reduced amounts of MAU2 in the chromatin fraction, suggesting the involvement of NIPBL-MAU2 interactions in the transcriptional regulation mediated by NIPBL.

The absence of MAU2 in the chromatin fraction of clone ex3mut (with no apparent effect on SCS) is interesting. Because *MAU2* expression was increased, the reduction of MAU2 was caused by the loss of MAU2 protein in the cells. MAU2 lacks a nuclear localization sequence (9); therefore, the inability of NIPBL to bind to MAU2 could result in a substantial reduction of nuclear MAU2. Our observation that MAU2 is absent from the nucleus of 293FT clone ex3mut and HCT116 clone ex3mut1 further supports the role of NIPBL-MAU2 binding in facilitating the nuclear localization of MAU2.

Although *MAU2* germline mutations have not been specifically associated with any human genetic disorders, *MAU2* missense or nonsense mutations have been observed to occur at rates similar to those in *NIPBL*, suggesting the importance of MAU2 function in human development (30). Elimination of MAU2 has been shown to cause developmental defects similar to those of *NIPBL* deletion in *Xenopus* and conditional knockout mouse models, suggesting that similar transcriptional disturbance underlies developmental defects during embryogenesis (9,26). Further studies are warranted to deepen our understanding of the transcriptional regulation mechanisms of the NIPBL-MAU2 heterodimer.

We did not observe a correlation between altered RAD21 genome-wide binding and transcriptional changes. Because reduced binding of RAD21 was a genome-wide phenomenon, expression abnormalities may have been triggered in genes predisposed for transcriptional alteration. We found that gene expression level may serve as such a predisposing factor. More specifically, genes with low WT expression levels tended to be overexpressed upon *NIPBL* inactivation both in *NIPBL* genome-edited cell lines and CdLS samples. We previously demonstrated the association between the degree of altered RNA polymerase II distribution and gene expression alterations, as well as the involvement RNA polymerase II pausing/elongation abnormalities, in CdLS (14). Therefore, the effect of transcriptional pausing/elongation defects may differ between downregulated and upregulated genes.

Another possible explanation for the lack of correlation between RAD21 binding alteration and transcriptional disturbance is that the transcriptional alteration may be a direct consequence of *NIPBL* mutations rather than alterations in RAD21 genome-wide binding. In yeast, the Scc2-Scc4 heterodimer (NIPBL-MAU2 homolog) is required for maintaining nucleosome-free regions (31). Disruption of this function may be a possible mechanism of the transcriptional disturbance observed upon deletion of the N-terminal region of NIPBL.

From a molecular genetics standpoint, it is important to recognize the possibility that a truncated protein can be created with the use of an alternative translational initiation site, even with the presence of a gene frameshift mutation. Likewise, when gene knockout mouse/cell lines are generated, careful evaluation to detect the potential presence of alternative gene transcripts from the same genomic locus should be performed.

In summary, we used the CRIPSR/Cas9 gene editing system to further elucidate the functional roles of NIPBL. Full-length NIPBL does not appear to be required for the maintenance of sister chromatid cohesion during cell division, and the N-terminus of NIPBL plays a central role in transcriptional regulation via interactions with MAU2. The resemblance of the mutations introduced into the genome-edited cell lines to those observed in individuals with CdLS and the various roles performed by cohesin provide evidence that the disruption of NIPBL-mediated transcriptional regulation constitutes the disease mechanism of CdLS.

## ACKNOWLEDGEMENTS

The authors thank Ms. Naoko Yokota, Ms. Keiko Nakagawa, and Mrs. Yuri Nakagawa for their assistance.

