## Supplemental file for "Dissecting the functions of NIPBL using genome editing: The importance of the N-terminus of NIPBL in transcriptional regulation"

**SUPPLEMENTARY TABLES AND FIGURES**

**Supplementary Table S1.** Next-generation sequencing statistics including RNA-seq, nascent RNA-seq, and RAD21 ChIP-seq statistics

**Supplementary Table S2.** Differentially expressed genes in 293FT cell line clone ex3mut

**Supplementary Figure S1.** Sanger sequencing chromatogram of *NIPBL* genome-edited cell lines. (**A**) 293FT cell line clone ex3mut. (**B**) 293FT clone ex10mut. (**C**) HCT116 clone ex3mut1. (**D**) HCT116 clone ex3mut2. (**E**) HCT116 clone ex10mut1. (**F**) HCT116 clone ex10mut2.

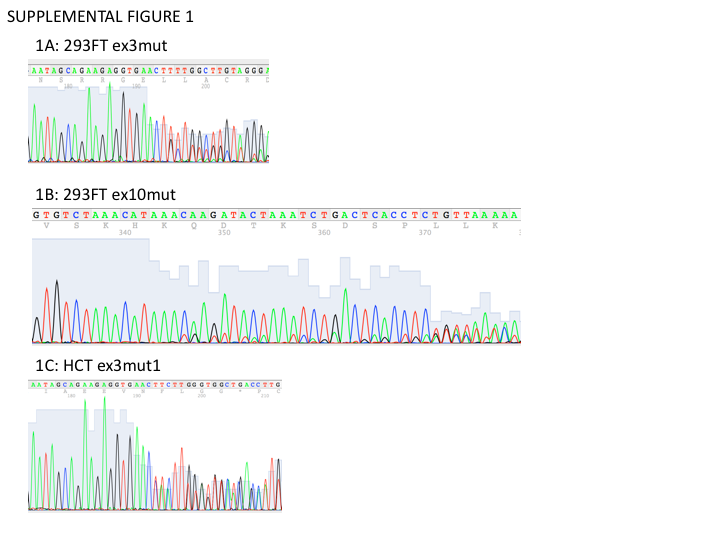

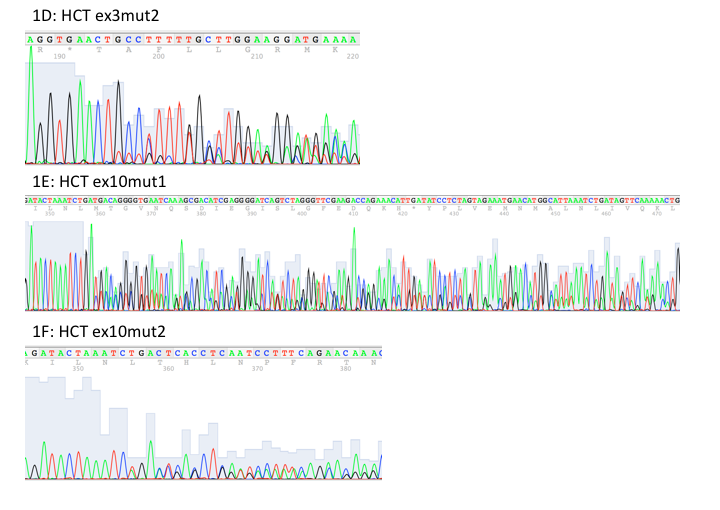

**Supplementary Figure S2.** HCT116 *NIPBL* mutant cell lines. (**A**) Western blotting results for HCT116 cell line clone ex3mut1 shows the absence of full-length NIPBL as assessed with antibodies directed against the N-terminus and C-terminus of NIPBL. The presence of truncated NIPBL is shown. (**B**) RNA-seq results for the *NIPBL* locus, indicating the presence of a transcript missing only a few base pairs targeted by *NIPBL* gRNA. (**C**) Western blotting for MAU2 and SMC1A in the total cellular lysate, soluble fraction (Sup), and chromatin fraction (Chr) of the HCT116 cell lines. MAU2 was not detected in the chromatin fraction of clone ex3mut1. The amount of MAU2 was also reduced in clones ex3mut2 and ex10mut1.

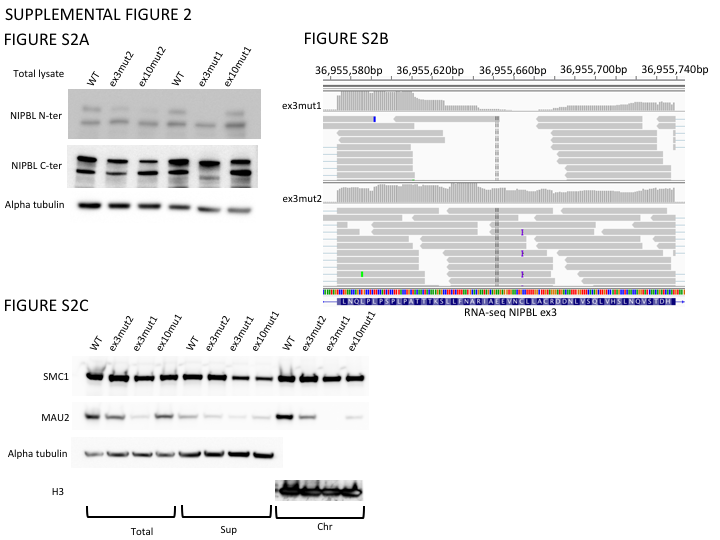

**Supplementary Figure S3.** Confirmation of the *NIPBL* isoform lacking exon 10. Reverse transcription-PCR using cDNA and primers specific for exon 9 and for exon 12 yielded an ~500-bp PCR product corresponding to a section of mRNA lacking exon 10.

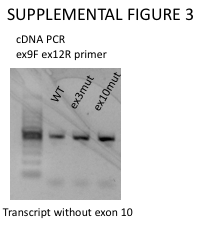

**Supplementary Figure S4.** Predicted TSS of 293FT cell line clone ex3mut.

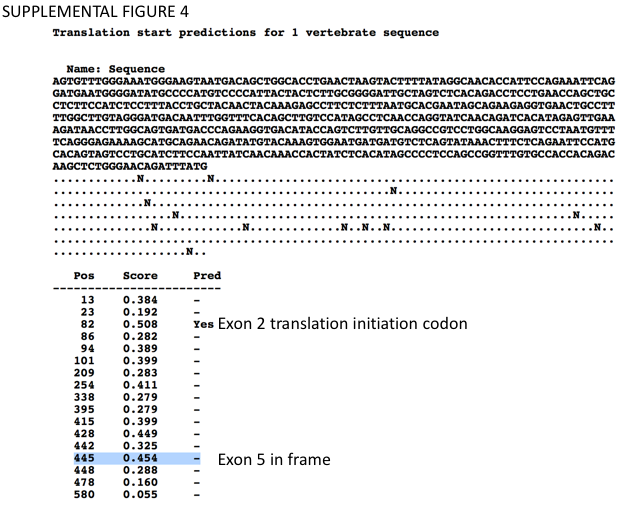

**Supplementary Figure S5.** Fluorescence-activated cell sorting results. The cell-cycle distribution was similar among WT 293FT cells and clones ex3mut and ex10mut.

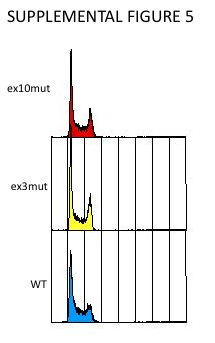

**Supplementary Figure S6.** Correlation between RNA-seq and DRB-nascent RNA-seq. The *x* axis depicts the log fold expression level difference between clone ex3mut and WT as assessed with RNA-seq. The *y* axis depicts the log fold expression level difference between clone ex3mut and WT as assessed with DRB-nascent RNA seq.

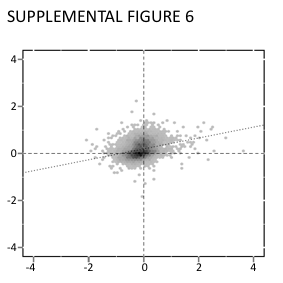
